# Mutations in *HUA2* restore flowering in the Arabidopsis *trehalose 6-phosphate synthase1* (*tps1*) mutant

**DOI:** 10.1101/2024.01.31.578264

**Authors:** Liping Zeng, Vasiliki Zacharaki, Yanwei Wang, Markus Schmid

## Abstract

Plant growth and development are regulated by many factors, including carbohydrate availability and signaling. Trehalose 6-phosphate (T6P), which is synthesized by TREHALOSE-6-PHOSPHATE SYNTHASE 1 (TPS1), is positively correlated with and functions as a signal that informs the cell about the carbohydrate status. Mutations in *TPS1* negatively affect the growth and development of *Arabidopsis thaliana* and complete loss-of-function alleles are embryo lethal, which can be overcome using inducible expression of *TPS1* (*GVG::TPS1*) during embryogenesis. Using EMS mutagenesis in combination with genome re-sequencing we have identified several alleles in the floral regulator *HUA2* that restore flowering and embryogenesis in *tps1-2 GVG::TPS1*. Genetic analyses using a *HUA2* T-DNA insertion allele, *hua2-4*, confirmed this finding. RNA-seq analyses demonstrated that *hua2-4* has widespread effects on the *tps1-2 GVG::TPS1* transcriptome, including key genes and pathways involved in regulating flowering. Higher order mutants combining *tps1-2 GVG::TPS1* and *hua2-4* with alleles in the key flowering time regulators *FLOWERING LOCUS T* (*FT*), *SUPPRESSOR OF OVEREXPRESSION OF CONSTANS 1* (*SOC1*), and *FLOWERING LOCUS C* (*FLC*) were constructed to analyze the role of *HUA2* during floral transition in *tps1-2* in more detail. Taken together, our findings demonstrate that loss of *HUA2* can restore flowering and embryogenesis in *tps1-2 GVG::TPS1* in part through activation of *FT*, with contributions of the upstream regulators *SOC1* and *FLC*. Interestingly, we found that mutation of *FLC* is sufficient to induce flowering in *tps1-2 GVG::TPS1*. Furthermore, we observed that mutations in *HUA2* modulate carbohydrate signaling and that this regulation might contribute to flowering in *hua2-4 tps1-2 GVG::TPS1*.

## Introduction

Plants have evolved intricate signaling mechanisms that enable them to monitor a wide range of environmental and endogenous cues and adjust their physiology, growth, and development accordingly. Adjustments occur more or less constantly, but developmental phase transitions such as germination, the switch from juvenile to adult growth, or the induction of flowering and reproductive development are under particularly stringent control.

In *Arabidopsis thaliana*, the floral transition is controlled by environmental factors including exposure to prolonged periods of cold (vernalization), ambient temperature, day length (photoperiod), light quality, and endogenous signals such as plant age, diverse hormones including gibberellic acid (GA), and carbohydrate signaling (Srikanth and Schmid, 2011; Romera-Branchat et al., 2014; Cho et al., 2017). Eventually, these signaling pathways converge on and regulate the expression of key floral integrator genes such as *FLOWERING LOCUS T* (*FT*) and *SUPPRESSOR OF OVEREXPRESSION OF CONSTANS 1* (*SOC1*) (Kardailsky et al., 1999; Moon et al., 2005; Kobayashi and Weigel, 2007; Turck et al., 2008; Lee and Lee, 2010; Jung et al., 2012). *FT* is induced in response to permissive photoperiod in the leaf vasculature where it is also translated. The FT protein is then transported via the phloem to the shoot apical meristem (SAM) where it interacts with the bZIP transcription factor FD and 14-3-3 proteins to form the florigen activation complex (FAC) (Abe et al., 2005; Wigge et al., 2005; Mathieu et al., 2007; Taoka et al., 2011; Collani et al., 2019). In contrast, *SOC1* is induced and acts largely at the SAM, both downstream and in parallel to *FT* (Yoo et al., 2005; Lee and Lee, 2010). Eventually, these factors induce flower meristem identity genes such as *LEAFY* (*LFY*) and *APETALA1* (*AP1*) at the SAM, thus completing the floral transition (Weigel and Nilsson, 1995; Liljegren et al., 1999; Blázquez and Weigel, 2000).

Apart from photoperiod, carbohydrate signaling has been shown to be necessary for *FT* expression (Wahl et al., 2013). Sucrose is the major product of photosynthesis and most common transport-sugar. However, rather than measuring sucrose concentration directly, plants employ trehalose 6-phosphate (T6P) as a readout and signal of sucrose availability (Goddijn and van Dun, 1999; Lunn et al., 2006; Martins et al., 2013; Yadav et al., 2014; Figueroa and Lunn, 2016). T6P is the intermediate of trehalose synthesis. It is synthesized from glucose 6-phosphate and uridine diphosphate glucose by TREHALOSE 6-PHOSPHATE SYNTHASE (TPS) and subsequently dephosphorylated by TREHALOSE 6-PHOSPHATE PHOSPHATASE (TPP) (Cabib and Leloir, 1958).

In *Arabidopsis thaliana*, there are 11 *TPS* genes (*AtTPS1*–*AtTPS11*), which can be divided into two subclades, class I and class II, and 10 TPP genes (*TPPA*–*TPPJ*) (Leyman et al., 2001; Lunn, 2007; Vandesteene et al., 2012). Of the class I *TPS* genes (*AtTPS1*–*AtTPS4*), only *AtTPS1*, *AtTPS2*, and *AtTPS4* have demonstrable catalytic activity, whereas *AtTPS3* harbors a premature translational stop codon and is likely a pseudogene (Blázquez et al., 1998; Van Dijck et al., 2002; Lunn, 2007; Delorge et al., 2015). Class II *TPS* genes (*AtTPS5*–*AtTPS11*), for which no TPS activity, has been detected, which have been reported to participate in cell size regulation, thermotolerance, and cold and salt resistance, but the underlying molecular mechanisms remain largely unclear (Chary et al., 2008; Ramon et al., 2009; Singh et al., 2011; Tian et al., 2019; Van Leene et al., 2022). The main T6P synthase in *Arabidopsis thaliana* is TPS1. *TPS1* loss-of-function mutants are embryonic lethal (Eastmond et al., 2002), but homozygous *tps1-2* mutants could be established by dexamethasone-inducible expression of *TPS1* (*GVG::TPS1*) during embryogenesis (van Dijken et al., 2004). Interestingly, the resulting homozygous *tps1-2 GVG:TPS1* plants flower extremely late compared to wild type under both short- and long-day conditions. At the molecular level, late flowering of *tps1-2 GVG::TPS1* has been attributed to the combined misregulation of key flowering time genes. In particular, *tps1-2 GVG::TPS1* mutant plants fail to induce *FT* in leaves even under permissive photoperiod. In addition, *MIR156* and its targets, the *SQUAMOSA PROMOTER BINDING PROTEIN LIKE* (*SPL*) genes, which together constitute the age pathway, are also misregulated in *tps1-2 GVG::TPS1* (Wahl et al., 2013). Nevertheless, many questions regarding the regulation of plant growth and development by the T6P pathway remain open.

In an EMS suppressor screen, we have recently reported dozens of mutations that partially restored flowering and seed set in *tps1-2 GVG::TPS1*, including several alleles in *SNF1 KINASE HOMOLOG 10* (*KIN10*) and *HOMOLOG OF YEAST SUCROSE NONFERMENTING 4* (*SNF4*), two subunits of *Arabidopsis thaliana SNF1-Related Kinase 1* (*SnRK1*) (Jung et al., 2012; Zacharaki et al., 2022), an evolutionary conserved regulator of cellular energy homeostasis.

Here, we identified several new alleles in *HUA2* (*At5g23150*) that partially rescue the *tps1-2 GVG::TPS1* phenotype. Mutations in *HUA2* were originally identified in a genetic screen as enhancers of the *AGAMOUS* (*AG*) allele *ag-4* (Chen and Meyerowitz, 1999). In addition, *HUA2* has also been reported to affect shoot morphology and function as a repressor of flowering (Doyle et al., 2005; Wang et al., 2007). At the molecular level, HUA2 has been suggested to function as a putative transcription factor but has also been implicated in RNA processing (Cheng et al., 2003). We show that three different EMS-induced point mutations in *HUA2* restore flowering in *tps1-2 GVG::TPS1* and verify this finding using a previously described T-DNA insertion allele, *hua2-4*. RNA-seq analyses revealed widespread effects of *hua2-4* on the *tps1 GVG::TPS1* transcriptome, including activation of flower integrator genes such as *SOC1* and *AGAMOUS-LIKE 24* (*AGL24*). Genetic analyses demonstrated that induction of flowering in *tps1-2 GVG::TPS1* required functional *FT*. Furthermore, we observed that loss of *FLOWERING LOCUS C* (*FLC*) is sufficient to induce flowering in *tps1-2 GVG::TPS1*. Interestingly, *hua2-4* also attenuated the induction of known SnRK1 target genes in response to carbon starvation. Taken together, our results identify mutations in *HUA2* as suppressors of the non-flowering phenotype of *tps1-2 GVG::TPS1* and provide insights into the underlying genetic and molecular pathways.

## Results

### Mutations in *hua2* restore flowering in *tps1-2 GVG::TPS1*

To identify novel components of the T6P pathway, we recently conducted a suppressor screen in which the non-flowering *tps1-2 GVG::TPS1* mutant was subjected to ethyl methane sulfonate (EMS) mutagenesis. In total, 106 M2 mutant plants in which flowering and seed set was at least partially restored were isolated, and EMS-induced SNPs were identified by whole genome sequencing in a subset of 65 mutants (Zacharaki et al., 2022). To identify additional candidate suppressor genes in which SNPs were overrepresented, we expanded this list to 92 by sequencing the genomes of another 27 mutants (Table S1).

Analysis of these 92 genome sequences for genes with multiple independent EMS-induced mutations identified three SNPs in the coding sequence of *HUA2* (*AT5G23150*) (Table S2, S3). The three alleles result in non-synonymous amino acid substitutions, namely A983T, P455S, and R902C. We refer to these new EMS-induced suppressor lines as *hua2-11* (line #8-1-1), *hua2-12* (line #233-14-1), and *hua2-13* (line #164-9-1), respectively (Fig. 1A). The polymorphism R902C resides at the C-terminal end of the HUA2 CID motif (RNA Pol-II C-terminal domain (CTD) interaction domain). The *hua2-11* (line #8-1-1) allele was also detected in two additional suppressor lines, #57-2-1 and #30-34 (Table S2, S3). As these three lines share most EMS-induced SNPs genome-wide, we assume these lines originate from the same parental plant.

**Figure 1.**
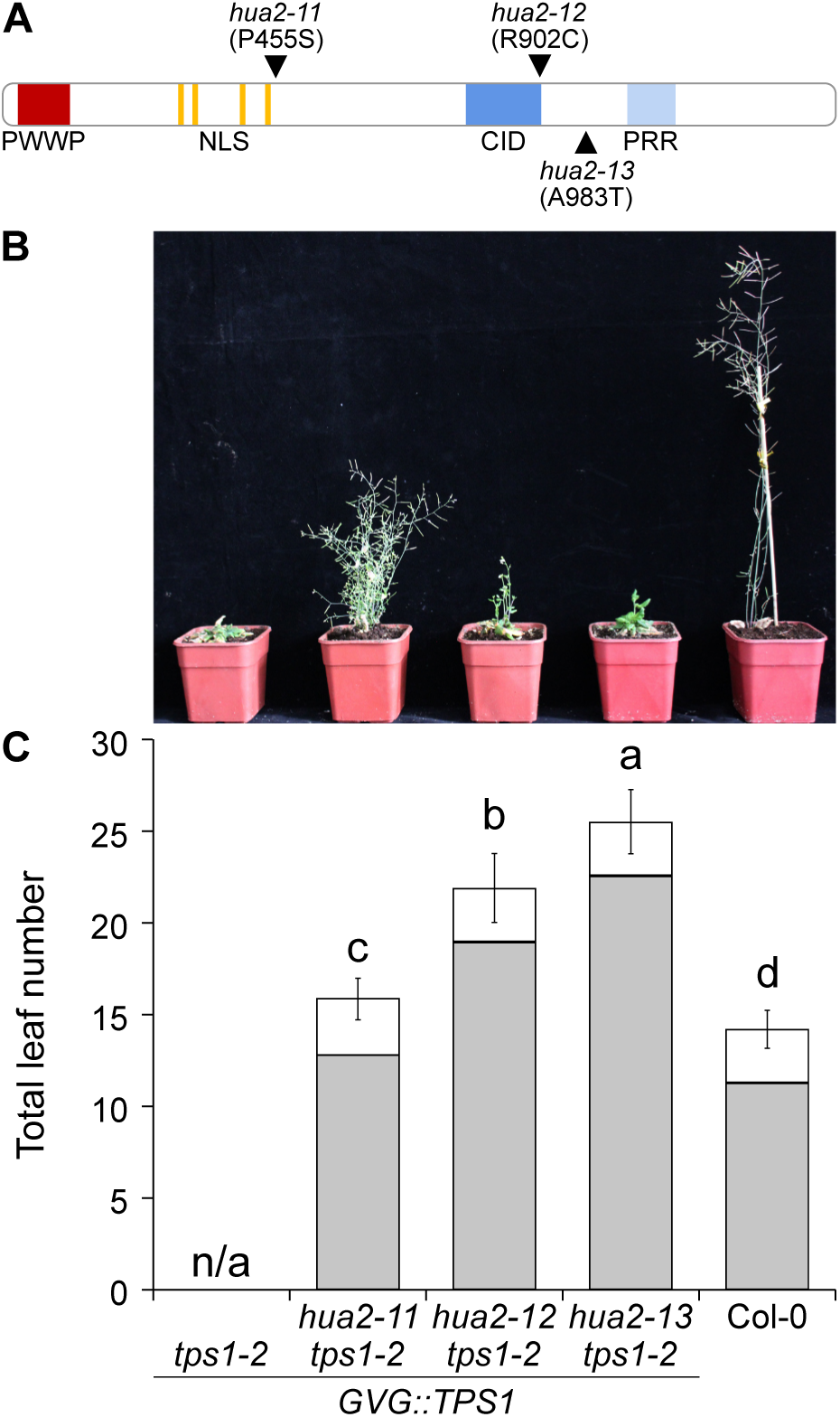
EMS-induced mutations in *HUA2* induce flowering in *tps1-2 GVG::TPS1* background. **A**) Schematic drawing of HUA2 indicating the position and the amino acid changes caused by the EMS-induced mutations *hua2-11* (P455S), *hua2-12* (R902C), and *hua2-13* (A983T). **B**) Phenotype of 9-week-old *tps1-2 GVG::TPS1*, *hua2-11 tps1-2 GVG::TPS1, hua2-12 tps1-2 GVG::TPS1, hua2-13 tps1-2 GVG::TPS1* and wild-type Col-0 *plants* grown in LD at 23°C. **C**) Flowering time of genotypes is given as total leaf number (rosette (grey); cauline leaves (white)) determined after bolting. Error bars represent the standard deviation. ANOVA Tukey’s multiple comparisons test was applied, and letters represent the statistical differences among genotypes (*P < 0.001*).

Importantly, flowering was restored in all three *hua2* alleles, even though all three mutant lines produced substantially more leaves before making the transition to flowering than Col-0 control plants (Fig. 1B, C). The flowering time of *hua2-11* was 32.15 days, whereas *hua2-12* and *hua2-13* flowered after 46.5 and 50.9 days, respectively, compared to Col-0, which flowered after 25.2 days. Thus, the three mutants form an allelic series with *hua2-11* being the strongest and *hua2-13* being the weakest allele. As *HUA2* has previously been implicated in flowering time regulation and has been shown to regulate the expression of a group of MADS-box transcription factors known to form a floral repressive complex in *Arabidopsis thaliana* (Doyle et al., 2005; Wang et al., 2007; Lee et al., 2013; Posé et al., 2013; Jali et al., 2014; Yan et al., 2016), we considered mutations in this gene as likely to be causal for the restoration of flowering in the *tps1-2 GVG::TPS1* suppressor lines.

Since the three *hua2* alleles described above were generated through EMS mutagenesis, it is possible that other independent mutations not linked to *HUA2* could be involved in partially rescuing the *tps1-2 GVG::TPS1* phenotype. To confirm that mutations in *HUA2* are causal for the suppression of the *tps1-2* non-flowering phenotype, we crossed *tps1-2 GVG::TPS1* with *hua2-4*, a previously described *hua2* loss-of-function mutant that carries a T-DNA insertion in the 2^nd^ intron (Fig. 2A) (Doyle et al., 2005). Of the F2 plants homozygous for the *tps1-2* mutations, only those approx. 25% that were homozygous for the *hua2-4* T-DNA insertion flowered without application of dexamethasone. Similar to *hua2-11 tps1-2 GVG::TPS1* (Fig. 1B,C), *hua2-4 tps1-2 GVG::TPS1* double mutants displayed a bushy shoot phenotype and were moderately late flowering (Fig. 2B,C). Taken together, our findings confirm that recessive mutations in *HUA2* are responsible for the induction of flowering in *tps1-2 GVG::TPS1*. Our findings also suggest that *HUA2* normally functions by repressing flowering either directly or indirectly through the promotion of floral repressors.

**Figure 2.**
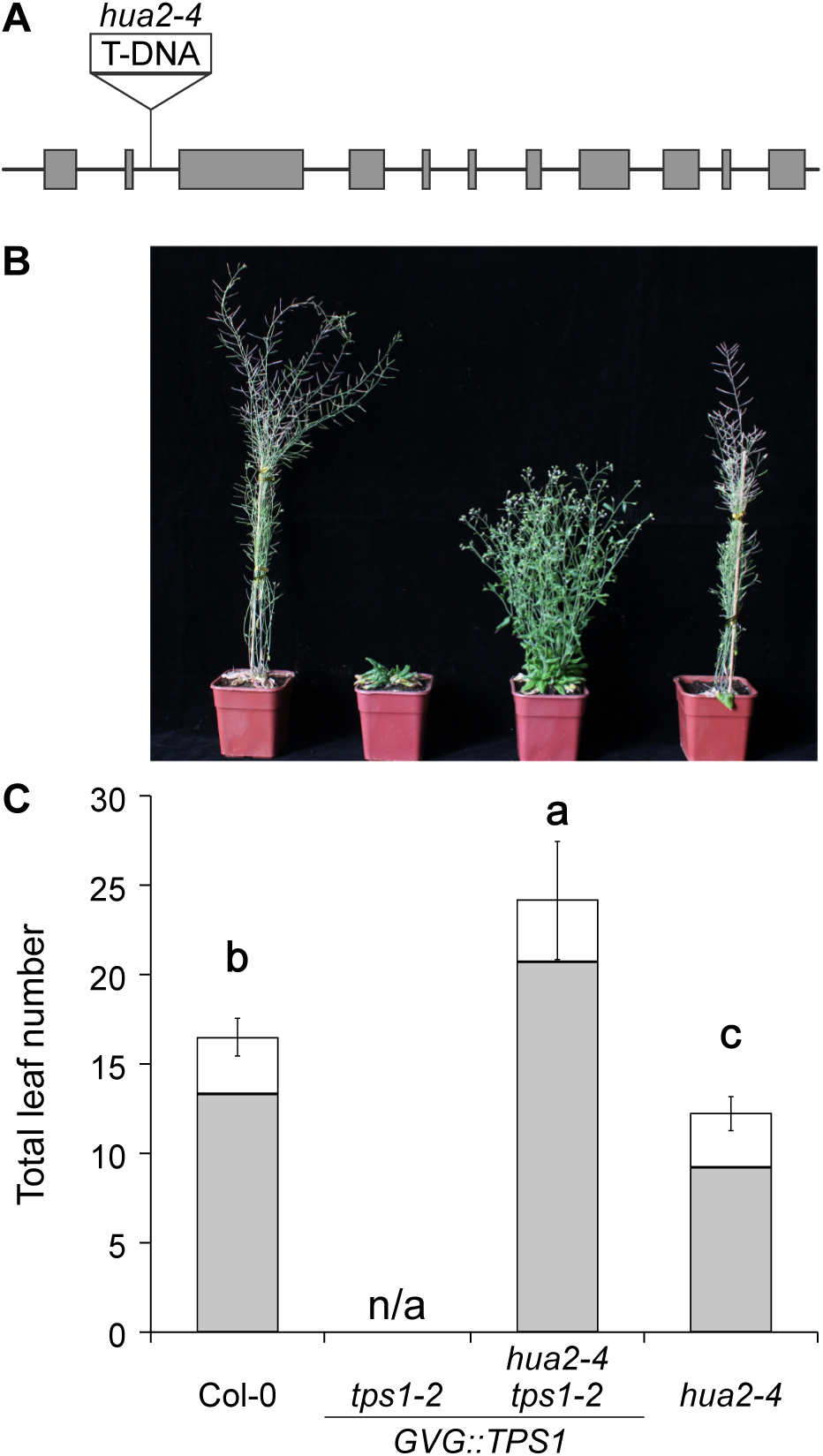
A T-DNA insertion in *HUA2* partially rescues the flowering time phenotype of *tps1-2 GVG::TPS1*. **A**) Schematic drawing of the *HUA2* locus indicating the position of the T-DNA insertion (SALK_032281C) in the 2^nd^ intron in *hua2-4*. **B-C**) Phenotypic analysis (**B**) and flowering time(**C**) of 9-week-old wild-type Col-0, *tps1-2 GVG::TPS1*, *hua2-4 tps1-2 GVG::TPS1* and *hua2-4* plants grown in LD at 23°C. Flowering time was scored as total leaf number (rosette (grey) and cauline leaves (white)) after bolting. Error bars represent the standard deviation. ANOVA Tukey’s multiple comparisons test was applied, and letters represent the statistical differences among genotypes (*P < 0.001*).

### *hua2-4* has widespread effects on the *tps1-2 GVG::TPS1* transcriptome

To identify possible downstream targets of *HUA2* whose misexpression might explain the induction of flowering in the suppressor mutant, we performed RNA-seq analysis in leaves of 21-d-old *tps1-2 GVG::TPS1* plants, *tps1-2 GVG::TPS1* plants treated with dexamethasone, and the *hua2-4 tps1-2 GVG::TPS1* double mutant. Plants were grown under long days (16 h light, 8 h dark) in the presence or absence of dexamethasone and samples were collected at ZT4 (Zeitgeber time 4, means 4 h after lights on). Genes that were differentially expressed in three independent replicates per genotype and treatment were identified using Cuffdiff.

We observed that dexamethasone treatment significantly affected the expression of 9600 genes in *tps1-2 GVG::TPS1*. Of these, 4830 and 4770 genes were upregulated and downregulated, respectively (Fig. 3A). In contrast, mutation of *hua2* affected the expression of only 2066 genes, of which 988 and 1078 genes were upregulated and downregulated in *hua2-4 tps1-2 GVG::TPS1*, respectively (Fig. 3A). In total our RNA-seq analysis identified 1437 genes that are differentially expressed in *tps1-2 GVG::TPS1* in response to dexamethasone application and the *hua2-4* mutation. Importantly, *HUA2* expression is not changed in *tps1-2 GVG::TPS1* in response to dexamethasone application, suggesting that *hua2* might induce flowering largely by activating a pathway not normally regulated by the T6P pathway (Fig. S1).

**Figure 3.**
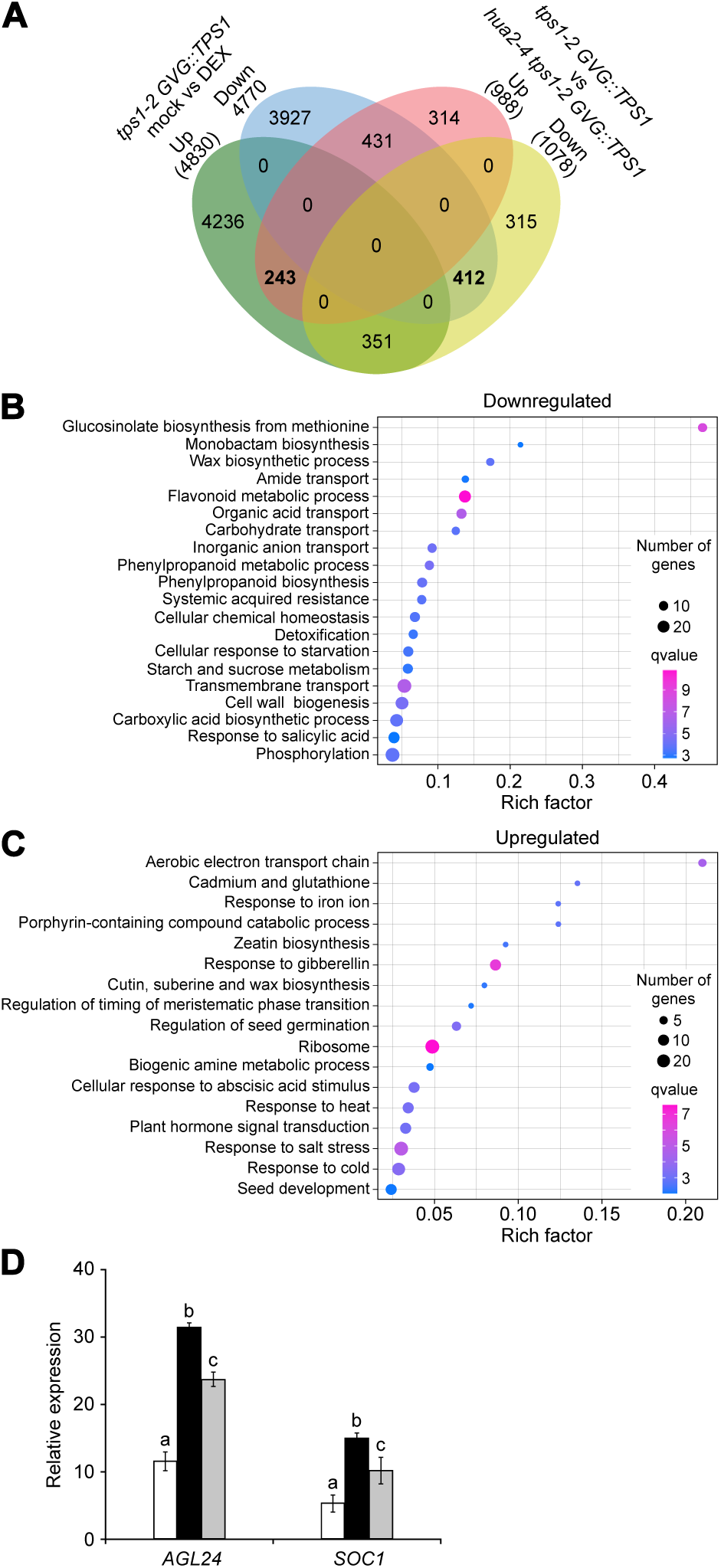
Characterization of *hua2-4 tps1-2 GVG::TPS1* transcriptome. **A**) 4-way Venn diagram of genes that are differentially expressed in *tps1-2 GVG::TPS1* in response to dexamethasone treatment and/or differentially expressed in *hua2-4 tps1-2 GVG::TPS1* when compared to *tps1-2 GVG::TPS1*. **B**) GO analysis of 412 genes downregulated in *tps1-2 GVG::TPS1* in response to dexamethasone treatment and in *hua2-4 tps1-2 GVG::TPS1*. **C)** GO analysis of 243 genes upregulated in *tps1-2 GVG::TPS1* in response to dexamethasone treatment and in *hua2-4 tps1-2 GVG::TPS1.* **D**) Relative expression of *AGL24* and *SOC1* in *tps1-2 GVG::TPS1* (white), *tps1-2 GVG::TPS1* treated with dexamethasone (black), and *hua2-4 tps1-2 GVG::TPS1* (grey). *AGL24* and *SOC1* are significantly differentially expressed. Error bars indicate the standard deviation. ANOVA Tukey’s multiple comparisons test was applied, and letters represent the statistical differences among genotypes (*P < 0.001*).

Since both, dexamethasone application and mutations in *hua2* can induce flowering in *tps1-2 GVG::TPS1,* we next searched for genes that were repressed or induced in response to either treatment. We identified 412 genes that were downregulated in *tps1-2 GVG::TPS1* in response to dexamethasone application and mutations in *hua2* (Fig. 3A). Gene ontology (GO) analysis revealed that among others, processes such as flavonoid metabolism (GO:0009812), carbohydrate transport (GO:0008643), and starvation response (GO:0009267) were significantly enriched, which is in line with the well-established role of *TPS1* in remodeling carbohydrate metabolism (Fig. 3B; Table S4).

In addition, we identified 243 genes that were induced in response to dexamethasone and in *hua2-4 tps1-2 GVG::TPS1*. Among these genes, GO categories related to the response to gibberellin (GO:0009739) and the regulation of timing of meristematic phase transition (GO:0048506) are of particular interest as they are directly linked to the transition to flowering (Fig. 3C; Table S5). Importantly, among the genes induced in *tps1-2 GVG::TPS1* by dexamethasone and *hua2* were *SUPPRESSOR OF OVEREXPRESSION OF CONSTANS 1* (*SOC1*) and *AGAMOUS-LIKE 24* (*AGL24*), two MADS-domain transcription factors known to promote the transition to flowering (Fig. 3D; Table S6). In contrast, other known flowering time regulators such as *CONSTANS* (*CO*), *FT*, and *TWIN SISTER OF FT* (*TSF*) are either hardly detectable (Fig. S2A), possibly because of the collection time of the RNA-seq samples at ZT 4 or did not change significantly in *hua2* and in response to dexamethasone treatment (Fig. S2B). In summary, our transcriptome analysis identified several downstream genes and pathways whose misregulation could contribute to the induction of flowering in *tps1-2 GVG::TPS1* in response to dexamethasone application or loss of *hua2* (Fig. S2; Table S6).

### Induction of flowering of *tps1-2 GVG::TPS1* by *hua2-4* requires *FT*

To test whether *SOC1*, which we found to be differentially expressed in response to dexamethasone application or in *hua2-4 tps1-2 GVG::TPS1*, is a major target of *HUA2* in the regulation of flowering time in *tps1-2 GVG::TPS1* we constructed the *soc1-2 hua2-4 tps1-2 GVG::TPS1* triple mutant. We observed that the triple mutant flowered only moderately later than the *hua2-4 tps1-2 GVG::TPS1* double mutant (Fig. 4A,B). This indicates that even though *SOC1* is significantly induced in our RNA-seq experiment in *hua2-4 tps1-2 GVG::TPS1* (Fig. 3D; Table S6) and in RT-qPCR experiments (Fig. 4C), *SOC1* is largely dispensable for the induction of flowering *in tps1-2 GVG::TPS1* by loss of *hua2*.

**Figure 4.**
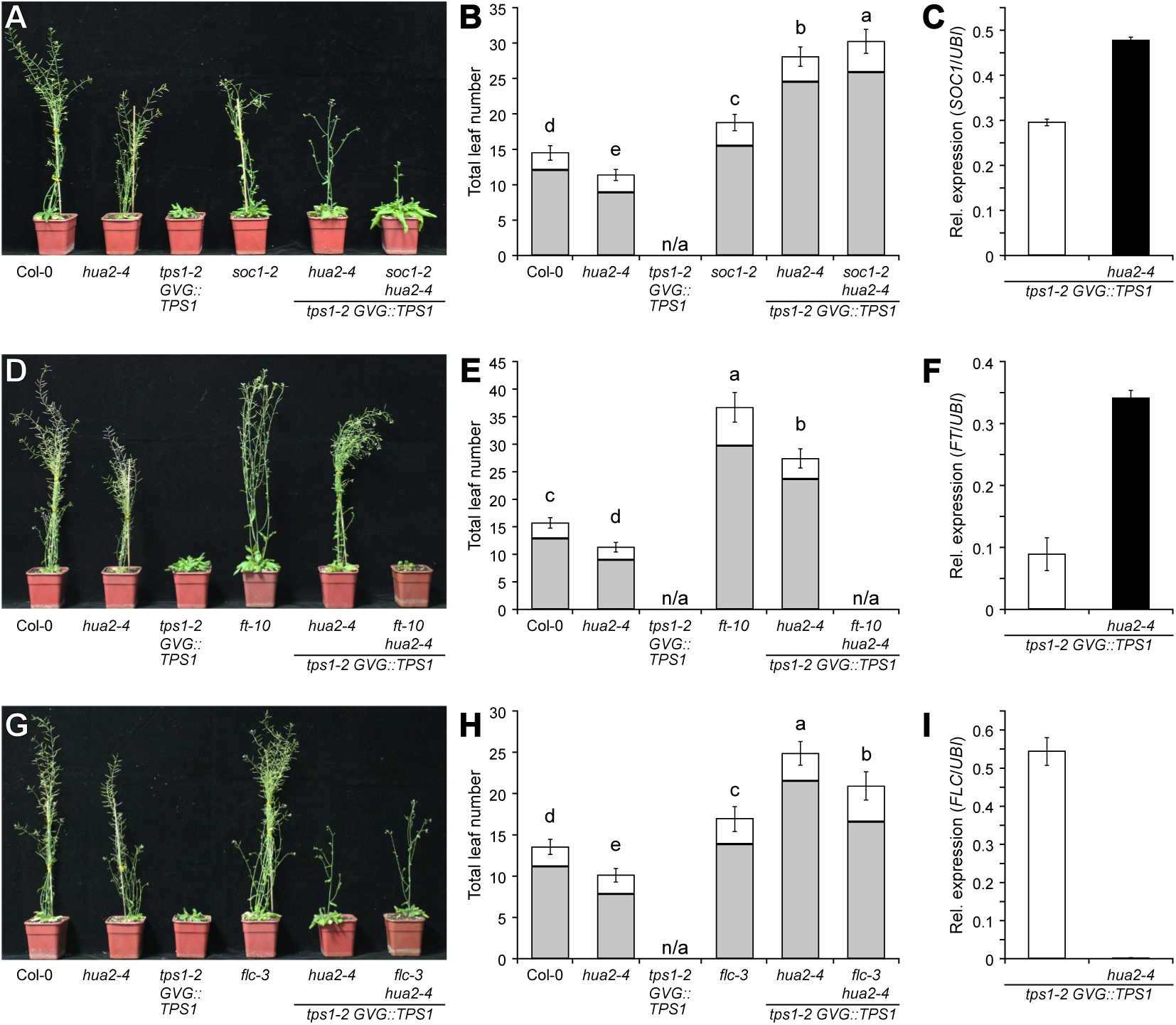
Genetic interactions between *tps1-2*, *hua2-4*, and floral regulators *SOC1*, *FT*, and *FLC*. **A-B**) Phenotypes (**A**) and flowering time (**B**) of Col-0, *hua2-4*, *tps-2 GVG::TPS1*, and *soc1-2* mutant combinations. **D-E**) Phenotypes (**D**) and flowering time (**E**) of Col-0, *hua2-4*, *tps-2 GVG::TPS1*, and *ft-10* mutant combinations. **G-H**) Phenotypes (**G**) and flowering time (**H**) of Col-0, *hua2-4*, *tps-2 GVG::TPS1*, and *flc-3* mutant combinations. Flowering time (**B**, **E**, **H**) was scored as total leaf number (rosette (grey) and cauline leaves (white)) after bolting. **C, F, I**) Relative expression of *SOC1* (**C**), *FT* (**F**), and *FLC* (**I**) in *tps1-2 GVG::TPS1* and *hua2-4 tps-2 GVG::TPS1*. Gene expression was determined by RT-qPCR at the end of the long day (ZT 16). Error bars represent the standard deviation. ANOVA Tukey’s multiple comparisons test was applied, and letters represent the statistical differences among genotypes (*P < 0.001*).

SOC1 is known to act partially upstream of the flowering time integrator gene and florigen FT. We, therefore, decided to test if induction of flowering in *tps1-2 GVG::TPS1* by *hua2-4* required functional *FT*. Interestingly, mutation of *FT* completely abolished the effect of *hua2-4* on flowering of *tps1-2 GVG::TPS1* and the *ft-10 hua2-4 tps1-2 GVG::TPS1* triple mutant failed to flower even after four months of growth in inductive long-day conditions (Fig. 4D,E). In line with this observation, we detected increased expression of *FT* at the end of the long day (ZT 16) in the *hua2-4 tps1-2 GVG::TPS1* double mutant when compared to *tps1-2 GVG::TPS1* (Fig. 4F). It is interesting to note that *FT* expression was barely detectable at ZT 4 according to our RNA-seq analysis (Fig. S2A), which is in agreement with the diurnal expression pattern reported for *FT* (Kobayashi et al., 1999). Taken together, our genetic and molecular analyses indicate that *hua2-4* induces flowering of *tps1-2 GVG::TPS1* in part through activation of *FT*, with minor contributions of the upstream regulators *SOC1*.

### Loss of *FLC* induces flowering in *tps1-2 GVG::TPS1*

*HUA2* has previously been reported to regulate flowering at least in part by regulating the expression of floral repressors of the MADS-domain transcription factor family, including *FLOWERING LOCUS C* (*FLC*) and *FLOWERING LOCUS M* (*FLM*) (Doyle et al., 2005). To test if *hua2-4* induces flowering in *tps-2 GVG::TPS1* through these repressors we constructed the *flc-3 hua2-4 tps1-2 GVG::TPS1* triple mutant. We found that this triple mutant flowered moderately earlier than *hua2-4 tps1-2 GVG::TPS1* (Fig. 4G,H). In agreement with these findings, RT-qPCR analysis failed to detect *FLC* expression in the *hua2-4 tps1-2 GVG::TPS1* mutant, whereas *FLC* expression was readily detectable by RT-qPCR in *tps1-2 GVG::TPS1* (Fig. 4I).

Furthermore, we found that the expression of *FLC* was significantly upregulated in 18-day-old *tps1-2 GVG::TPS1* seedlings when compared to Col-0 in publicly available RNA-seq data (Zacharaki et al., 2022) (Fig. 5A). This prompted us to test loss off *FLC* on its own might be sufficient to suppress the non-flowering phenotype of *tps1-2 GVG::TPS1*. Indeed, we observed that *flc-*3 alone is capable of inducing flowering in the otherwise non-flowering *tps1-2 GVG::TPS1* mutant background, even though the *flc-3 tps1-2 GVG::TPS1* double mutant flowered significantly later than wild-type and *flc-3* (Fig. 5B,C). These findings suggest that the failure of *tps1-2 GVG::TPS1* to flower could in part be due to *FLC*, possibly in conjunction with other MADS-box repressors such as *MADS AFFECTING FLOWERING 5* (*MAF5*), the expression of which was also elevated in *tps1-2 GVG::TPS1* (Fig. 5A). In contrast, expression of *HUA2* was not changed in *tps1-2 GVG::TPS1* when compared to Col-0 according to publicly available RNA-seq data (Fig. S3).

**Figure 5.**
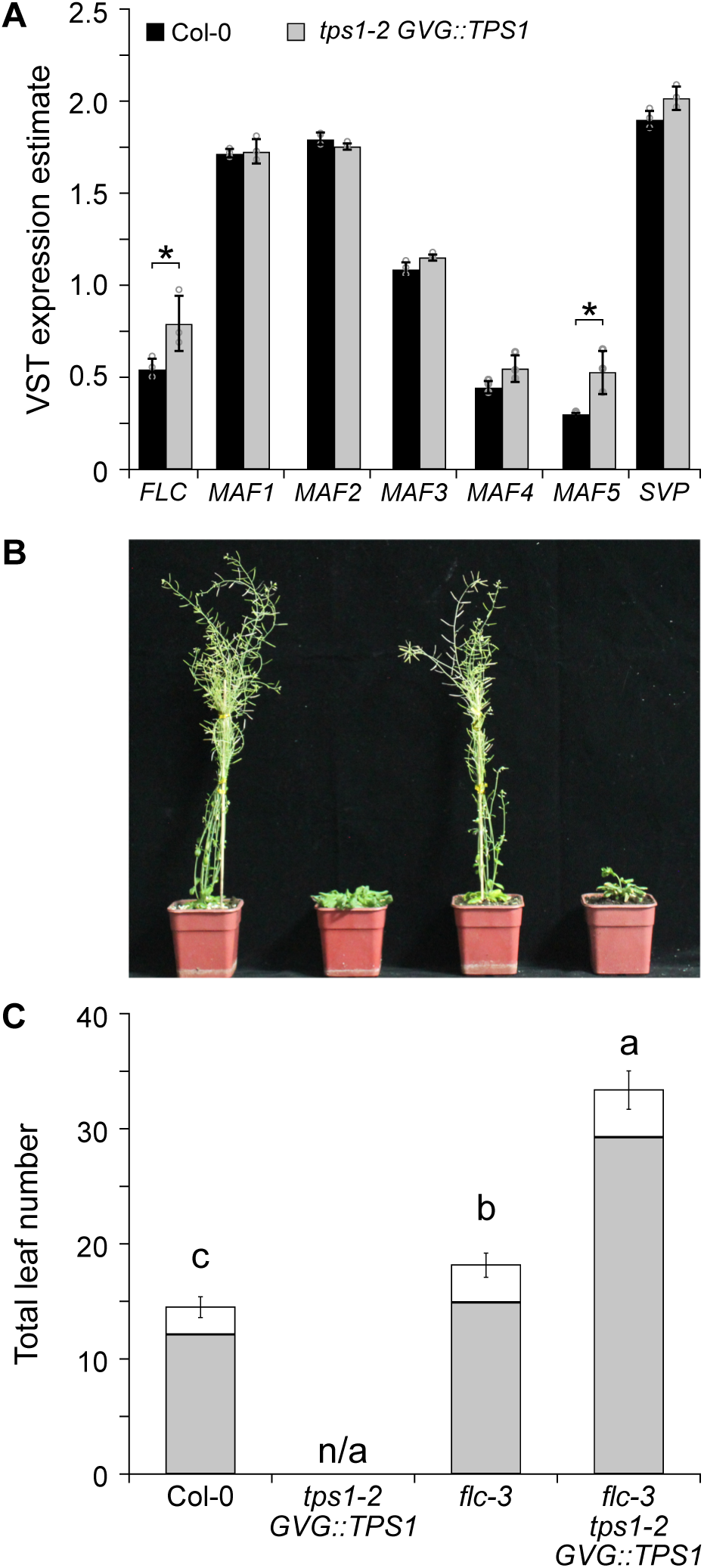
Loss of *FLC* rescues the non-flowering phenotype of *tps1-2 GVG::TPS1*. **A**) VST expression estimates for MADS-box floral repressors in 18-day-old plants. RNA-seq expression data retrieved from Zacharaki et al., 2022. Columns indicate mean VST expression estimates as implemented in DEseq2 calculated from three individual biological replicates per genotype. Circles indicate expression estimates for individual biological replicates. Asterisks indicate differential gene expression with a statistical significance of *Padj < 0.01*. **B-C**) Phenotypes (**B**) and total leaf number (**C**) of Col-0, *tps1-2 GVG::TPS1*, *flc-3*, and *flc-3 tps1-2 GVG::TPS1 double mutant.* Flowering time was scored as total leaf number (rosette (grey) and cauline leaves (white)) after bolting. Error bars represent the standard deviation. ANOVA Tukey’s multiple comparisons test was applied, and letters represent the statistical differences among genotypes (*P < 0.001*).

### *hua2-4* attenuates carbon starvation responses

The above data indicate that mutations in *HUA2* bypass the requirement for *TPS1* to induce flowering by reducing expression of MADS-box floral repressors and ultimately inducing floral integrator genes such as *FT* and *SOC1*. However, carbohydrate signaling has been shown to also indirectly regulate phase transitions, including flowering, in *A. thaliana* (Corbesier et al., 1998; Gibson, 2005; Xing et al., 2015; Wang et al., 2020). In part, this response is mediated by SnRK1, which in response to stress conditions such as extended darkness phosphorylates a range of proteins, including several C- and S1-class bZIP transcription factors. Activation of these transcription factors by SnRK1 induces expression of stress response genes, including *SENESCENCE5* (*SEN5*) and *DARK INDUCED6*/*ASPARAGINE SYNTHASE1* (*DIN6*/*ASN1*), which can be used as a proxy for SnRK1 activity (Delatte et al., 2011; Dietrich et al., 2011; Mair et al., 2015). To test if loss of *HUA2* might affect flowering also more indirectly by modulating cellular energy responses, we analyzed the expression of *SEN5* and *DIN6*. Interestingly, we found that induction of *SEN5* and *DIN6* in response to extended night was strongly attenuated in *hua2-4* (Fig. 6A, B) similar to what we had previously observed in mutants affected in SnRK1 subunits (Zacharaki et al., 2022). This finding indicates that mutations in *HUA2* might modulate carbohydrate signaling more directly and that this regulation might contribute to the induction of flowering in *hua2-4 tps1-2 GVG::TPS1*. In agreement with this hypothesis, we found that expression of *SEN5* and *DIN6* was even further attenuated in three independent *hua2-4 tps1-2 GVG::TPS1* lines (Fig. 6A,B).

**Figure 6.**
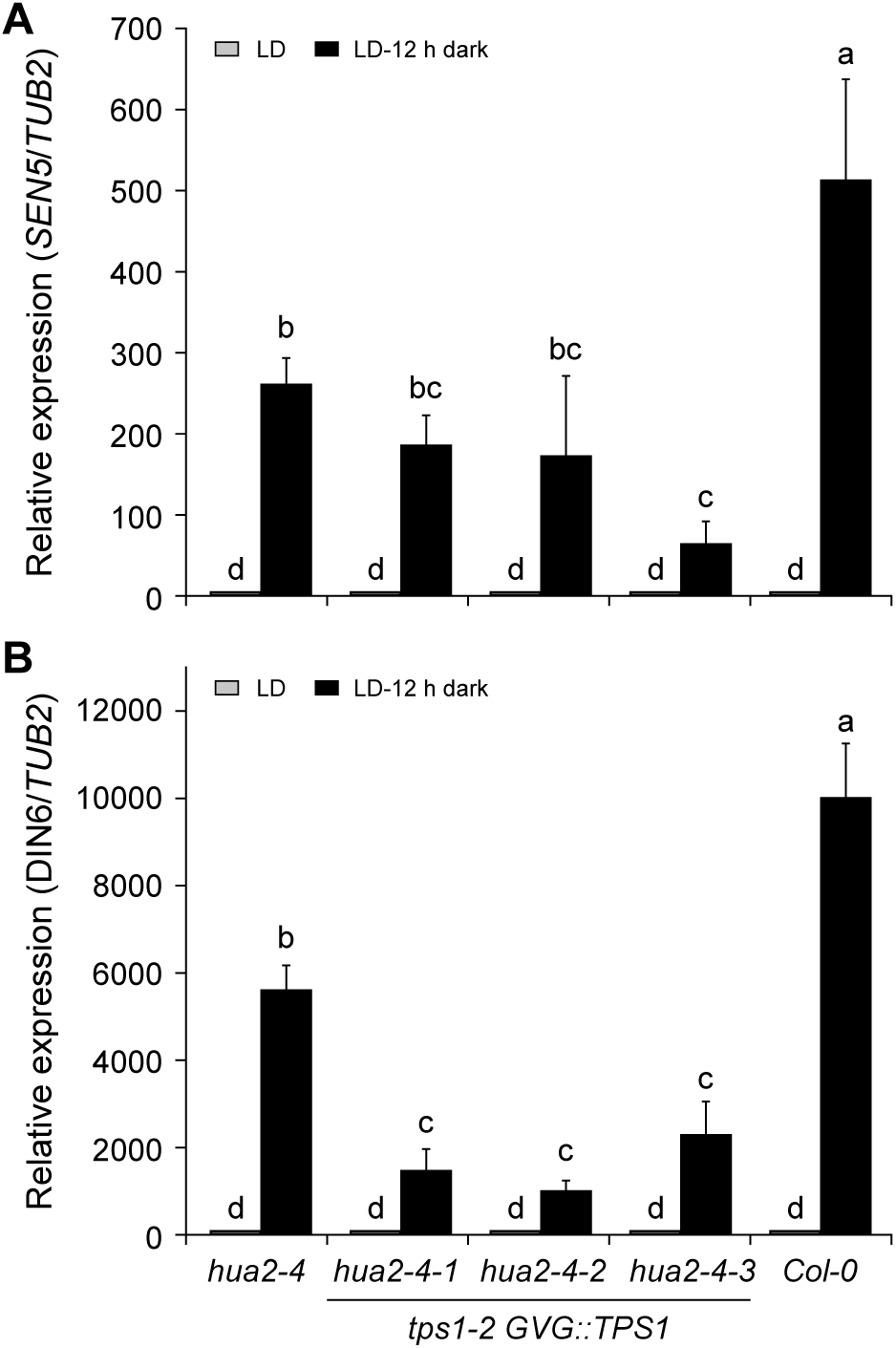
Expression of SnRK1 target genes *SEN5* and *DIN6* in *hua2-4* and *hua2-4 tps1-2 GVG::TPS1* double mutant. **A-B**) Induction of *SEN5* (**A**) and *DIN6* (**B**) in response to extended night is attenuated in 14-day-old of *hua2-4* single mutant and three independent lines of the *hua2-4 tps1-2 GVG::TPS1* double mutant. Plants were grown for 14 days in LD (grey) before being exposed to a single extended night (12h additional darkness; black). LD, long days. Error bars represent the standard deviation. ANOVA Tukey’s multiple comparisons test was applied, and letters represent the statistical differences among genotypes (*P < 0.001*).

## Discussion

*Arabidopsis thaliana HUA2* has been reported to play a crucial role in various aspects of plant growth and development. *HUA2* was initially identified as an enhancer of the *AGAMOUS* (*AG*) allele *ag-4* (Chen and Meyerowitz, 1999). Later, *HUA2* was found to also play a role as a repressor of flowering (Doyle et al., 2005; Wang et al., 2007). At the molecular level, *HUA2* has been suggested to function as a putative transcription factor but has also been implicated in RNA processing (Cheng et al., 2003). *HUA2* is expressed throughout the whole plant growth period (Chen and Meyerowitz, 1999), indicating the importance and widespread effects on plant growth. Here, our study showed that loss of *HUA2* can partially restore flowering and embryogenesis in *tps1-2 GVG::TPS1*.

It is interesting to note that in our EMS suppressor screen, we did not identify mutations in any of the *HUA2-like* genes, *HULK1, HULK2,* and *HUL3*, present in *A. thaliana* (Jali et al., 2014). One possible explanation is that our genetic screen might not have been saturated or that *HUA2*-like genes were missed due to the relatively low sequencing coverage. However, we believe this to be rather unlikely given that our approach has recovered multiple alleles in *HUA2* (this study) as well as two SnRK1 subunits (Zacharaki et al., 2022). Furthermore, flowering time is unaffected in the *hua2-like* single mutants and *hulk2 hulk3* double mutants have been shown to be late flowering (Jali et al., 2014). Thus, it seems unlikely that mutation in any of the *HUA2-like* genes would suppress the non-flowering phenotype of *tps1-2 GVG::TPS1*.

*HUA2* has been reported to exert its function in part by regulating the expression of MADS-box transcription factors (Doyle et al., 2005), named after *MINICHROMOSOME MAINTENANCE 1* (*MCM1*) in yeast, *AGAMOUS* (*AG*) in Arabidopsis, *DEFICIENS* (*DEF*) in Antirrhinum, and serum response factor (SRF) in humans. MADS-BOX domain transcription factors contribute to all major aspects of the life of land plants, such as female gametophyte, floral organ identity, seed development, and flowering time control (Portereiko et al., 2006; Colombo et al., 2008; Koo et al., 2010; Lee et al., 2013; Posé et al., 2013). In this context, it is interesting to note that our transcriptome and genetic analysis identified several MADS-box transcription factors to be misregulated in *tps1-2 GVG::TPS1*. In particular, the well-known floral repressors *FLOWERING LOCUS C* (*FLC*) and *MADS AFFECTING FLOWERING5* (*MAF5*) were found to be induced in *tps1-2 GVG::TPS1* compared to Col-0 (Fig. 5A). Moreover, loss of *FLC* was sufficient to induce flowering in *tps1-2 GVG::TPS1* (Fig. 5B,C), suggesting that these floral repressors are partially responsible for the non-flowering phenotype of *tps1-2 GVG::TPS1*. Our transcriptome analyses further identified two MADS-box transcription factors, *SUPPRESSOR OF OVEREXPRESSION OF CONSTANS 1* (*SOC1*) and *AGAMOUS-LIKE 24* (*AGL24*), both known to promote flowering in Arabidopsis, to be upregulated in *hua2-4 tps1-2 GVG::TPS1*.

The molecular mechanism by which *HUA2* regulates the expression of these MADS-box flowering time regulators is currently unclear. However, since HUA2 localizes to the nucleus, it seems possible that HUA2 is directly involved in regulating the expression of these genes. For example, *HUA2* could (directly) promote the expression of *FLC*, which has previously been shown to directly bind to and repress the expression of *FT* and *SOC1* (Chen and Meyerowitz, 1999; Doyle et al., 2005; Deng et al., 2011). In such a scenario, the increased expression of *FT*, *SOC1*, and *AGL24* in *hua2-4 tps1-2 GVG::TPS1* would be the result of reduced expression of floral repressors such as *FLC* and *MAF5.* However, the regulation of flowering is a very complex process full of intricate feedback loops and *HUA2* might regulate *SOC1* and *AGL24* directly, rather than indirectly. In this context, it is interesting to note that a nonfunctional *hua2* allele may compensate for the loss of *FLC* in L*er* accession (Lemus et al., 2023). Alternatively, HUA2 might affect the expression of these important flowering time genes through interaction with RNA Pol-II via its CID domain, which is affected by the *hua2-13* alleles (R902C). Interestingly, polymorphisms resulting in amino acid substitutions in natural accessions of *A. thaliana* have been reported for R902 and A983, but not for P455 (The 1001 Genomes Consortium, 2016). Even though the molecular mechanisms underlying *HUA2* function remain elusive, our results confirm *HUA2* as a central regulator of flowering time in *Arabidopsis thaliana*.

We have previously identified mutations in two subunits of *SNF1-Related Kinase 1* (*SnRK1*), *KIN10* and *SNF4*, that partially restore flowering and seed set in *tps1-2 GVG::TPS1* (Zacharaki et al., 2022). Identification of these suppressor mutations was in line with the role of SnRK1 as a downstream regulator of the T6P pathway and other stresses. Antagonizing SnRK1 in the regulation of energy homeostasis in plants is target of rapamycin (TOR), the activity of which is inhibited under energy-limiting conditions (Baena-González and Hanson, 2017; Belda-Palazón et al., 2022). In contrast to mutations in *KIN10* and *SNF4*, mutations in *HUA2* appear, at first glance, to be bypass mutations that induce flowering independently of T6P signaling. However, and rather unexpectedly, we did observe that mutation of *HUA2* attenuated the induction of the carbon starvation markers *SEN5* and *DIN6* in response to extended night treatments (Fig. 6A, B), indicating that mutations in *HUA2* might modulate carbohydrate signaling more directly than anticipated. How exactly *HUA2* modulates carbon responses in Arabidopsis remains to be established. It is well-known that T6P signaling through *SnRK1* affects processes such as carbon starvation response, germination, flowering, and senescence in opposition to the TOR (target of rapamycin) pathway (Figueroa and Lunn, 2016; Baena-González and Lunn, 2020). The regulatory network controlling this central metabolic hub is still not fully understood and new players are constantly added. For example, it has recently been shown that class II TPS proteins are important negative regulators of *SnRK1* (Van Leene et al., 2022).

Regarding a possible role of HUA2 in integrating carbon responses, it is worth noting that flavonoid-related genes (GO:0009812) were downregulated in *tps1-2 GVG::TPS1* in response to dexamethasone application and the *hua2* mutant (Fig. 3B). This is interesting as HUA2 is known to promote anthocyanin accumulation (Ilk et al., 2015), whereas SnRK1 has been shown to repress sucrose-induced anthocyanin production (Li et al., 2014; Meng et al., 2018; Brouke et al., 2023). Thus, HUA2 might constitute an important hub in coordinating metabolic responses. However, as expression of *SnRK1* subunits is not affected in *hua2-4 tps1-2 GVG::TPS1* when compared to *tps1-2 GVG::TPS1* (Fig. S4), such a role would likely be indirect.

Understanding the interplay between energy metabolism, in particular SnRK1, TOR, and T6P signaling, and plant growth and development is of utmost importance for developing plants capable of withstanding future challenges. The suppressor mutants generated in the *tps1-2 GVG::TPS1* background comprise an important resource in our hunt for additional factors that, like *HUA2*, link energy metabolism to plant development.

## Material and methods

### Plant materials and growth conditions

All T-DNA insertion mutants and transgenic lines used in this work are in the Col-0 background. The *tps1-2 GVG::TPS1* line used in this work is referred to as ind-TPS1 #201 in the original publication (Dijken et al., 2004). The *hua2-4* (SALK_032281C) was obtained from NASC and the presence of the T-DNA insertion was confirmed by PCR (Table S5). *ft-10* (GABI-Kat: 290E08) was provided by Dr. Yi Zhang, Southern University of Science and Technology, *flc-3* (Kim et al., 2006) by Dr. Liangyu Liu, Capital Normal University, and *soc1-2* (Lee et al., 2000) by Dr. Jie Luo, Chinese Academy of Sciences. *tps1-2 GVG::TPS1 hua2-4* plants were generated by crossing and double homozygous mutants were identified by phenotyping and genotyping of F2 individuals. Higher order mutants were obtained by crossing *soc1-2*, *flc-3*, and *ft-10* mutants with the *tps1-2 GVG::TPS1 hua2-4* double mutant and homozygous triple mutants were identified in the F2 and F3 generation. All mutant genotypes were confirmed by PCR, see Table S7 for details. Plants were planted onto nutrient soil with normal water supply and grown under either long days (LD) with a photoperiod of 16 hours light at 22°C and 8 hours darkness at 20°C or in short days (SD) with a photoperiod of 8 hours light at 22°C and 16 hours darkness at 20°C. Flowering time are presented as average rosette leaf number, cauline leaf number, and total leaf number.

### Genome sequencing and analysis

Young leaves were used for DNA extraction for sequencing using the NovaSeq 6000 Sequencing platform (Novogene). Adapters and low-quality sequences of raw reads were trimmed using Trimmomatic (Bolger et al., 2014), and the clean reads were mapped to the reference genome of Col-0 using BWA-MEM (v0.7.15) (Cingolani et al., 2012). SNP calling was performed using Genome Analysis Toolkit 4 (GATK4; https://gatk.broadinstitute.org/hc/en-us) with default parameters. Variants were annotated using snpEff 4.3 (Li and Durbin, 2009) based on TAIR 10 annotation. Next, we identified the protein-coding genes with multiple non-redundant mutations and found three mutant lines harboring unique non-synonymous mutations in the *HUA2* gene. The method was inspired by our previous study that multiple EMS-induced mutants with unique mutation sites in the coding regions of *SnRK1* alpha subunit rescued the non-flowering phenotype of *tps1* (Zacharaki et al., 2022).

### Gene expression analysis by RNA-seq

For RNA-seq analyses, plants were grown on soil for 3 weeks in LD conditions. Leaves from 21-day-old *Arabidopsis thaliana* were collected, immediately snap-frozen and stored at −80 °C. Total RNA was extracted using RNAprep Pure Plant Plus Kit (Tiangen, China, DP441). RNA integrity was assessed using the RNA Nano 6000 Assay Kit on the Bioanalyzer 2100 system (Agilent Technologies, CA, USA). RNA-seq libraries were generated with three independent biological replicates and sequenced on the Illumina NovaSeq platform by Annoroad Gene Technology. The raw RNA-seq reads were quality trimmed by Trimmomatic (v 0.11.9) (Bolger et al., 2014). The qualified reads were mapped to TAIR10 version genome guided by gene annotation model using HISAT2 (v2.1.0) (Kim et al., 2015). The expression level for each gene was determined by StringTie (v1.3.4) (Pertea et al., 2016). The differential expressed genes were identified by DESeq2 (Love et al., 2014).

### RNA isolation and RT-qPCR data analysis

Total RNA was extracted from Arabidopsis seedlings using the RNA Isolation Kit (Tiangen, China, DP441) according to the manufacturer’s instructions. cDNA was synthesized from 3 µg total RNA in a 10 µl reaction volume using the RevertAid Premium First Strand cDNA Synthesis Kit (Fermentas, Thermo Fisher Scientific, Rochester, NY). Quantitative real-time PCR (qRT-PCR) was performed using TB Green™ Premix Ex Taq™ II (Takara, Dalian, China). Relative gene expression was calculated using the 2−ΔΔCt method (Rao et al., 2013). All analyses were repeated three times. The primer used for qRT-PCR are listed in Supplemental Tables S5.

### Accession numbers

Identifiers of key genes used in this study: *TPS1* (At1g78580), *HUA2* (AT5G23150), *SOC1* (AT2G45660), *FLC* (AT5G10140), *FT* (AT1G65480). RNA-seq data generated in this study have been deposited with NCBI under the BioProject PRJNA1005425.

## Data availability

The data and material that support the findings of this study are available from the corresponding author upon reasonable request.

## Author contributions

LZ and MS designed the experiments. LZ carried out the SNP detection and genetic analyses with input from VZ and MS. LZ carried out the gene expression analyses. LP and MS wrote the manuscript with contributions from all authors.

## Acknowledgments

The authors would like to thank Ruben M. Benstein and Vanessa Wahl for discussion and comments on the manuscript. This work was supported by grants from the National Natural Science Foundation of China (32071504 and 32371577) YW, the fundamental research funds for the central universities (BLX202170) to LZ, and the DFG (SPP1530: SCHM1560/8-1, 8-2) and the Swedish Research Council (2015-04617) to MS.

## Supplemental Material

**Supplemental Figure S1** Relative expression of *HUA2* in *tps1-2 GVG::TPS1* treated with dexamethasone or untreated.

**Supplemental Figure S2** Relative expression of important floral regulators in *tps1-2 GVG::TPS1*, *tps1-2 GVG::TPS1* treated with dexamethasone, and *hua2-4 tps1-2 GVG::TPS1*.

**Supplemental Figure S3** VST expression estimates for *HUA2* in 18-day-old plants.

**Supplemental Figure S4** Relative expression of SnRK1 subunits in *tps1-2 GVG::TPS1*, *tps1-2 GVG::TPS1* treated with dexamethasone, and *hua2-4 tps1-2 GVG::TPS1*.

**Supplemental Table S1** Number of SNPs identified in individual suppressor mutants.

**Supplemental Table S2** Number of SNPs identified in EMS suppressor lines carrying mutations in *HUA2*.

**Supplemental Table S3** EMS suppressor lines bearing non-synonymous mutations in *HUA2*.

**Supplemental Table S4** GO analysis of 412 genes downregulated in *tps1-2 GVG::TPS1* in response to dexamethasone application and in *hua2-4*.

**Supplemental Table S5** GO analysis of 243 genes induced in *tps1-2 GVG::TPS1* in response to dexamethasone application and in *hua2-4*.

**Supplemental Table S6** Expression of flowering time genes in *hua2-4 tps1-2 GVG::TPS1* and *tps1-2 GVG::TPS1*.

**Supplemental Table S7** List of oligonucleotides used in this study.

## References

Abe M, Kobayashi Y, Yamamoto S, Daimon Y, Yamaguchi A, Ikeda Y, Ichinoki H, Notaguchi M, Goto K, Araki T (2005) FD, a bZIP protein mediating signals from the floral pathway integrator FT at the shoot apex. Science 309: 1052–1056

Baena-González E, Hanson J (2017) Shaping plant development through the SnRK1-TOR metabolic regulators. Curr Opin Plant Biol 35: 152–157

Baena-González E, Lunn JE (2020) SnRK1 and trehalose 6-phosphate - two ancient pathways converge to regulate plant metabolism and growth. Curr Opin Plant Biol 55: 52–59

Belda-Palazón B, Costa M, Beeckman T, Rolland F, Baena-González E (2022) ABA represses TOR and root meristem activity through nuclear exit of the SnRK1 kinase. Proc Natl Acad Sci USA 119: 1–3

Blázquez MA, Santos E, Flores CL, Martínez-Zapater JM, Salinas J, Gancedo C (1998) Isolation and molecular characterization of the Arabidopsis TPS1 gene, encoding trehalose-6-phosphate synthase. Plant J 13: 685–689

Blázquez MA, Weigel D (2000) Integration of floral inductive signals in Arabidopsis. Nature 404: 889–892

Bolger AM, Lohse M, Usadel B (2014) Trimmomatic: a flexible trimmer for Illumina sequence data. Bioinformatics 30: 2114–2120

Broucke E, Dang TTV, Li Y, Hulsmans S, Van Leene J, De Jaeger G, Hwang I, Wim VdE, Rolland F (2023) SnRK1 inhibits anthocyanin biosynthesis through both transcriptional regulation and direct phosphorylation and dissociation of the MYB/bHLH/TTG1 MBW complex. Plant J 115: 1193–1213

Cabib E, Leloir LF (1958) THE BIOSYNTHESIS OF TREHALOSE PHOSPHATE. Journal of Biological Chemistry 231: 259–275

Chary SN, Hicks GR, Choi YG, Carter D, Raikhel NV (2008) Trehalose-6-phosphate synthase/phosphatase regulates cell shape and plant architecture in Arabidopsis. Plant Physiol 146: 97–107

Chen X, Meyerowitz EM (1999) HUA1 and HUA2 are two members of the floral homeotic AGAMOUS pathway. Mol Cell 3: 349–360

Cheng Y, Kato N, Wang W, Li J, Chen X (2003) Two RNA binding proteins, HEN4 and HUA1, act in the processing of AGAMOUS pre-mRNA in Arabidopsis thaliana. Dev Cell 4: 53–66

Cho LH, Yoon J, An G (2017) The control of flowering time by environmental factors. Plant J 90: 708–719

Cingolani P, Platts A, Wang le L, Coon M, Nguyen T, Wang L, Land SJ, Lu X, Ruden DM (2012) A program for annotating and predicting the effects of single nucleotide polymorphisms, SnpEff: SNPs in the genome of Drosophila melanogaster strain w1118; iso-2; iso-3. Fly (Austin) 6: 80–92

Collani S, Neumann M, Yant L, Schmid M (2019) FT Modulates Genome-Wide DNA-Binding of the bZIP Transcription Factor FD. Plant Physiol 180: 367–380

Colombo M, Masiero S, Vanzulli S, Lardelli P, Kater MM, Colombo L (2008) AGL23, a type I MADS-box gene that controls female gametophyte and embryo development in Arabidopsis. Plant J 54: 1037–1048

Corbesier L, Lejeune P, Bernier G (1998) The role of carbohydrates in the induction of flowering in Arabidopsis thaliana: comparison between the wild type and a starchless mutant. Planta 206: 131–137

Delatte TL, Sedijani P, Kondou Y, Matsui M, de Jong GJ, Somsen GW, Wiese-Klinkenberg A, Primavesi LF, Paul MJ, Schluepmann H (2011) Growth arrest by trehalose-6-phosphate: an astonishing case of primary metabolite control over growth by way of the SnRK1 signaling pathway. Plant Physiol 157: 160–174

Delorge I, Figueroa CM, Feil R, Lunn JE, Van Dijck P (2015) Trehalose-6-phosphate synthase 1 is not the only active TPS in Arabidopsis thaliana. Biochem J 466: 283–290

Deng W, Ying H, Helliwell CA, Taylor JM, Peacock WJ, Dennis ES (2011) FLOWERING LOCUS C (FLC) regulates development pathways throughout the life cycle of Arabidopsis. Proc Natl Acad Sci USA 108: 6680–6685

Dietrich K, Weltmeier F, Ehlert A, Weiste C, Stahl M, Harter K, Dröge-Laser W (2011) Heterodimers of the Arabidopsis transcription factors bZIP1 and bZIP53 reprogram amino acid metabolism during low energy stress. Plant Cell 23: 381–395

Doyle MR, Bizzell CM, Keller MR, Michaels SD, Song J, Noh YS, Amasino RM (2005) HUA2 is required for the expression of floral repressors in Arabidopsis thaliana. Plant J 41: 376–385

Eastmond PJ, van Dijken AJ, Spielman M, Kerr A, Tissier AF, Dickinson HG, Jones JD, Smeekens SC, Graham IA (2002) Trehalose-6-phosphate synthase 1, which catalyses the first step in trehalose synthesis, is essential for Arabidopsis embryo maturation. Plant J 29: 225–235

Figueroa CM, Lunn JE (2016) A Tale of Two Sugars: Trehalose 6-Phosphate and Sucrose. Plant Physiol 172: 7–27

Gibson SI (2005) Control of plant development and gene expression by sugar signaling. Curr Opin Plant Biol 8: 93–102

Goddijn OJ, van Dun K (1999) Trehalose metabolism in plants. Trends Plant Sci 4: 315–319

Ilk N, Ding J, Ihnatowicz A, Koornneef M, Reymond M (2014) Natural variation for anthocyanin accumulation under high-light and low-temperature stress is attributable to the ENHANCER OF AG-4 2 (HUA2) locus in combination with PRODUCTION OF ANTHOCYANIN PIGMENT1 (PAP1) and PAP2. New Phytol 206: 422–435

Jali SS, Rosloski SM, Janakirama P, Steffen JG, Zhurov V, Berleth T, Clark RM, Grbic V (2014) A plant-specific HUA2-LIKE (HULK) gene family in Arabidopsis thaliana is essential for development. Plant J 80: 242–254

Jung JH, Ju Y, Seo PJ, Lee JH, Park CM (2012) The SOC1-SPL module integrates photoperiod and gibberellic acid signals to control flowering time in Arabidopsis. Plant J 69: 577–588

Kardailsky I, Shukla VK, Ahn JH, Dagenais N, Christensen SK, Nguyen JT, Chory J, Harrison MJ, Weigel D (1999) Activation tagging of the floral inducer FT. Science 286: 1962–1965

Kim D, Langmead B, Salzberg SL (2015) HISAT: a fast spliced aligner with low memory requirements. Nat Methods 12: 357–360

Kobayashi Y, Kaya H, Goto K, Iwabuchi M, Araki T (1999) A pair of related genes with antagonistic roles in mediating flowering signals. Science 286: 1960–1962

Kobayashi Y, Weigel D (2007) Move on up, it’s time for change--mobile signals controlling photoperiod-dependent flowering. Genes Dev 21: 2371–2384

Koo SC, Bracko O, Park MS, Schwab R, Chun HJ, Park KM, Seo JS, Grbic V, Balasubramanian S, Schmid M, Godard F, Yun DJ, Lee SY, Cho MJ, Weigel D, Kim MC (2010) Control of lateral organ development and flowering time by the Arabidopsis thaliana MADS-box Gene AGAMOUS-LIKE6. Plant J 62: 807–816

Lee J, Lee I (2010) Regulation and function of SOC1, a flowering pathway integrator. J Exp Bot 61: 2247–2254

Lee JH, Ryu HS, Chung KS, Posé D, Kim S, Schmid M, Ahn JH (2013) Regulation of temperature-responsive flowering by MADS-box transcription factor repressors. Science 342: 628–632

Lemus T, Mason GA, Bubb KL, Alexandre CM, Queitsch C, Cuperus JT (2023) AGO1 and HSP90 buffer different genetic variants in *Arabidopsis thaliana*. Genetics 223: iyac163

Leyman B, Van Dijck P, Thevelein JM (2001) An unexpected plethora of trehalose biosynthesis genes in Arabidopsis thaliana. Trends Plant Sci 6: 510–513

Li H, Durbin R (2009) Fast and accurate short read alignment with Burrows-Wheeler transform. Bioinformatics 25: 1754–1760

Li Y, Van den Ende W, Rolland F (2014) Sucrose induction of anthocyanin biosynthesis is mediated by DELLA. Mol Plant 7: 570–572

Liljegren SJ, Gustafson-Brown C, Pinyopich A, Ditta GS, Yanofsky MF (1999) Interactions among APETALA1, LEAFY, and TERMINAL FLOWER1 specify meristem fate. Plant Cell 11: 1007–1018

Love MI, Huber W, Anders S (2014) Moderated estimation of fold change and dispersion for RNA-seq data with DESeq2. Genome Biology 15: 1–21

Lunn JE (2007) Gene families and evolution of trehalose metabolism in plants. Funct Plant Biol 34: 550–563

Lunn JE, Feil R, Hendriks JH, Gibon Y, Morcuende R, Osuna D, Scheible WR, Carillo P, Hajirezaei MR, Stitt M (2006) Sugar-induced increases in trehalose 6-phosphate are correlated with redox activation of ADPglucose pyrophosphorylase and higher rates of starch synthesis in Arabidopsis thaliana. Biochem J 397: 139–148

Mair A, Pedrotti L, Wurzinger B, Anrather D, Simeunovic A, Weiste C, Valerio C, Dietrich K, Kirchler T, Naegele T, Vicente Carbajosa J, Hanson J, Baena-Gonzalez E, Chaban C, Weckwerth W, Droege-Laser W, Teige M (2015) SnRK1-triggered switch of bZIP63 dimerization mediates the low-energy response in plants. Elife 4: 1–33

Martins MC, Hejazi M, Fettke J, Steup M, Feil R, Krause U, Arrivault S, Vosloh D, Figueroa CM, Ivakov A, Yadav UP, Piques M, Metzner D, Stitt M, Lunn JE (2013) Feedback inhibition of starch degradation in Arabidopsis leaves mediated by trehalose 6-phosphate. Plant Physiol 163: 1142–1163

Mathieu J, Warthmann N, Kuttner F, Schmid M (2007) Export of FT protein from phloem companion cells is sufficient for floral induction in Arabidopsis. Curr Biol 17: 1055–1060

Meng LS, Xu MK, Wan W, Yu F, Li C, Wang JY, Wei ZQ, Lv MJ, Cao XY, Li ZY, Jiang JH (2018) Sucrose signaling regulates anthocyanin biosynthesis through a MAPK cascade in Arabidopsis thaliana. Genetics 210: 607–619

Moon J, Lee H, Kim M, Lee I (2005) Analysis of flowering pathway integrators in Arabidopsis. Plant Cell Physiol 46: 292–299

Pertea M, Kim D, Pertea GM, Leek JT, Salzberg SL (2016) Transcript-level expression analysis of RNA-seq experiments with HISAT, StringTie and Ballgown. Nat Protoc 11: 1650–1667

Portereiko MF, Lloyd A, Steffen JG, Punwani JA, Otsuga D, Drews GN (2006) AGL80 is required for central cell and endosperm development in Arabidopsis. Plant Cell 18: 1862–1872

Posé D, Verhage L, Ott F, Yant L, Mathieu J, Angenent GC, Immink RG, Schmid M (2013) Temperature-dependent regulation of flowering by antagonistic FLM variants. Nature 503: 414–417

Ramon M, De Smet I, Vandesteene L, Naudts M, Leyman B, Van Dijck P, Rolland F, Beeckman T, Thevelein JM (2009) Extensive expression regulation and lack of heterologous enzymatic activity of the Class II trehalose metabolism proteins from Arabidopsis thaliana. Plant Cell Environ 32: 1015–1032

Romera-Branchat M, Andrés F, Coupland G (2014) Flowering responses to seasonal cues: what’s new? Curr Opin Plant Biol 21: 120–127

Singh V, Louis J, Ayre BG, Reese JC, Pegadaraju V, Shah J (2011) TREHALOSE PHOSPHATE SYNTHASE11-dependent trehalose metabolism promotes Arabidopsis thaliana defense against the phloem-feeding insect Myzus persicae. Plant J 67: 94–104

Srikanth A, Schmid M (2011) Regulation of flowering time: all roads lead to Rome. Cell Mol Life Sci 68: 2013–2037

Taoka K, Ohki I, Tsuji H, Furuita K, Hayashi K, Yanase T, Yamaguchi M, Nakashima C, Purwestri YA, Tamaki S, Ogaki Y, Shimada C, Nakagawa A, Kojima C, Shimamoto K (2011) 14-3-3 proteins act as intracellular receptors for rice Hd3a florigen. Nature 476: 332–335

The 1001 Genomes Consortium (2016) 1,135 genomes reveal the global pattern of polymophism in *Arabidopsis thaliana*. Cell 166: 481–491.

Tian L, Xie Z, Lu C, Hao X, Wu S, Huang Y, Li D, Chen L (2019) The trehalose-6-phosphate synthase TPS5 negatively regulates ABA signaling in Arabidopsis thaliana. Plant Cell Rep 38: 869–882

Turck F, Fornara F, Coupland G (2008) Regulation and identity of florigen: FLOWERING LOCUS T moves center stage. Annu Rev Plant Biol 59: 573–594

Van Dijck P, Mascorro-Gallardo JO, De Bus M, Royackers K, Iturriaga G, Thevelein JM (2002) Truncation of Arabidopsis thaliana and Selaginella lepidophylla trehalose-6-phosphate synthase unlocks high catalytic activity and supports high trehalose levels on expression in yeast. Biochem J 366: 63–71

van Dijken AJ, Schluepmann H, Smeekens SC (2004) Arabidopsis trehalose-6-phosphate synthase 1 is essential for normal vegetative growth and transition to flowering. Plant Physiol 135: 969–977

Van Leene J, Eeckhout D, Gadeyne A, Matthijs C, Han C, De Winne N, Persiau G, Van De Slijke E, Persyn F, Mertens T, Smagghe W, Crepin N, Broucke E, Van Damme D, Pleskot R, Rolland F, De Jaeger G (2022) Mapping of the plant SnRK1 kinase signalling network reveals a key regulatory role for the class II T6P synthase-like proteins. Nat Plants 8: 1245–1261

Vandesteene L, López-Galvis L, Vanneste K, Feil R, Maere S, Lammens W, Rolland F, Lunn JE, Avonce N, Beeckman T, Van Dijck P (2012) Expansive evolution of the trehalose-6-phosphate phosphatase gene family in Arabidopsis. Plant Physiol 160: 884–896

Wahl V, Ponnu J, Schlereth A, Arrivault S, Langenecker T, Franke A, Feil R, Lunn JE, Stitt M, Schmid M (2013) Regulation of flowering by trehalose-6-phosphate signaling in Arabidopsis thaliana. Science 339: 704–707

Wang M, Zang L, Jiao F, Perez-Garcia M-D, Ogé L, Hamama L, Le Gourrierec J, Sakr S, Chen J (2020) Sugar Signaling and Post-transcriptional Regulation in Plants: An Overlooked or an Emerging Topic? Front Plant Sci 11: 578096

Wang Q, Sajja U, Rosloski S, Humphrey T, Kim MC, Bomblies K, Weigel D, Grbic V (2007) HUA2 caused natural variation in shoot morphology of A. thaliana. Curr Biol 17: 1513–1519

Weigel D, Nilsson O (1995) A developmental switch sufficient for flower initiation in diverse plants. Nature 377: 495–500

Wigge PA, Kim MC, Jaeger KE, Busch W, Schmid M, Lohmann JU, Weigel D (2005) Integration of spatial and temporal information during floral induction in Arabidopsis. Science 309: 1056–1059

Xing LB, Zhang D, Li YM, Shen YW, Zhao CP, Ma JJ, An N, Han MY (2015) Transcription Profiles Reveal Sugar and Hormone Signaling Pathways Mediating Flower Induction in Apple (Malus domestica Borkh.). Plant Cell Physiol 56: 2052–2068

Yadav UP, Ivakov A, Feil R, Duan GY, Walther D, Giavalisco P, Piques M, Carillo P, Hubberten HM, Stitt M, Lunn JE (2014) The sucrose-trehalose 6-phosphate (Tre6P) nexus: specificity and mechanisms of sucrose signalling by Tre6P. J Exp Bot 65: 1051–1068

Yan W, Chen D, Kaufmann K (2016) Molecular mechanisms of floral organ specification by MADS domain proteins. Curr Opin Plant Biol 29: 154–162

Yoo SK, Chung KS, Kim J, Lee JH, Hong SM, Yoo SJ, Yoo SY, Lee JS, Ahn JH (2005) CONSTANS activates SUPPRESSOR OF OVEREXPRESSION OF CONSTANS 1 through FLOWERING LOCUS T to promote flowering in Arabidopsis. Plant Physiol 139: 770–778

Zacharaki V, Ponnu J, Crepin N, Langenecker T, Hagmann J, Skorzinski N, Musialak-Lange M, Wahl V, Rolland F, Schmid M (2022) Impaired KIN10 function restores developmental defects in the Arabidopsis trehalose 6-phosphate synthase1 (tps1) mutant. New Phytol 235: 220–233

