## Supplemental Figures 1-4 and Tables 1-3,6,7 for "Mutations in *HUA2* restore flowering in the Arabidopsis *trehalose 6-phosphate synthase1* (*tps1*) mutant"

### Supplementary Material

- –Supplementary Figure S1
- –Supplementary Figure S2
- –Supplementary Figure S3
- –Supplementary Figure S4
- –Supplementary Table S1
- –Supplementary Table S2
- –Supplementary Table S3
- –Supplementary Table S6
- –Supplementary Table S7

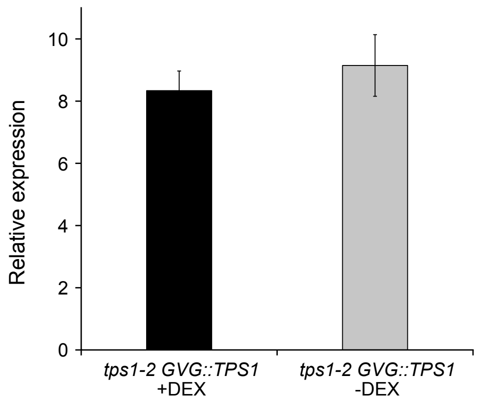


**Figure S1.** Relative expression of *HUA2* in *tps1-2 GVG::TPS1* treated with dexamethasone (black) or untreated (grey). Error bars indicate SD. ANOVA Tukey's multiple comparisons test was applied. No statistically significant difference in *HUA2* expression was detected.

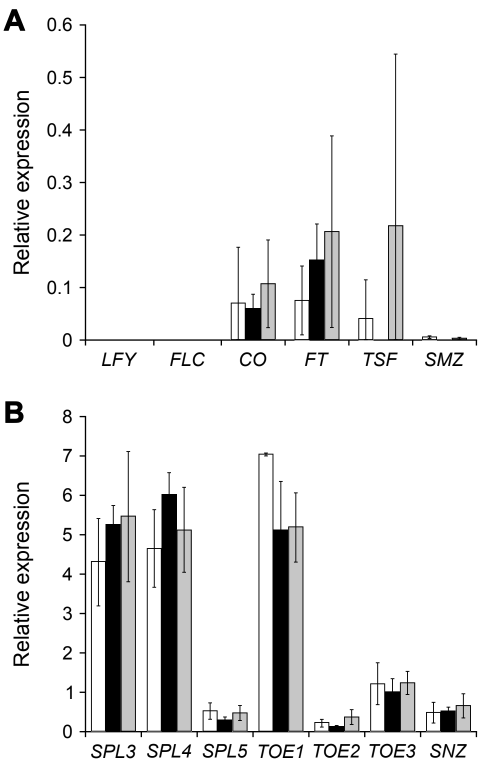


**Figure S2.** Relative expression of important floral regulators in *tps1-2 GVG::TPS1* (white), *tps1-2 GVG::TPS1* treated with dexamethasone (black), and *hua2-4 tps1-2 GVG::TPS1* (grey). (A) floral regulators are expressed at low levels (B) or not differentially expressed. Error bars indicate SD. ANOVA Tukey's multiple comparisons test was applied. No statistically significant differences in gene expression between genotypes were detected.

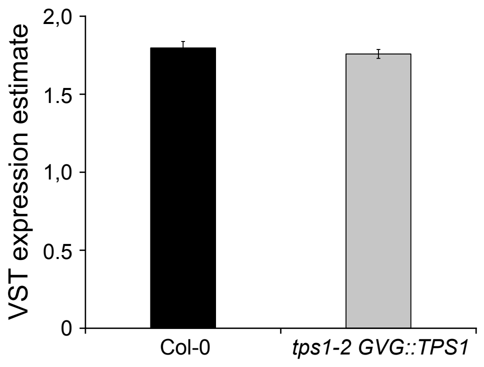


**Figure S3.** VST expression estimates for *HUA2* in 18-day-old plants. RNA-seq expression data retrieved from Zacharaki et al., 2022. Columns indicate mean VST expression estimates as implemented in DEseq2 calculated from three individual biological replicates per genotype. No statistically significant difference in *HUA2* expression was detected.

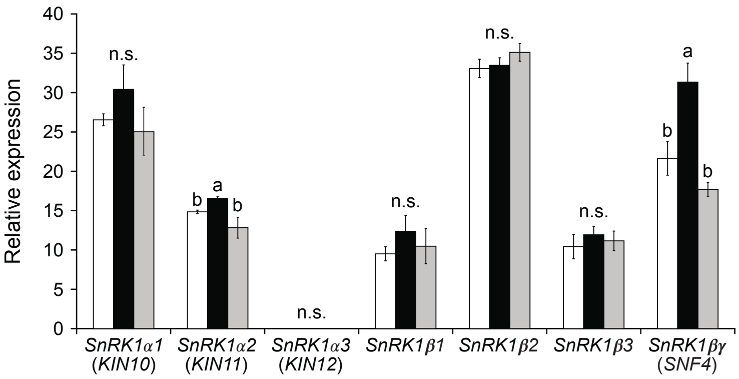


**Figure S4.** Relative expression of SnRK1 subunits in *tps1-2 GVG::TPS1* (white), *tps1-2 GVG::TPS1* treated with dexamethasone (black), and *hua2-4 tps1-2 GVG::TPS1* (grey). Error bars indicate SD. ANOVA Tukey's multiple comparisons test was applied.

**Table S1. Number of SNPs identified in individual suppressor mutants.**

| Nr | Line ID | SNPs | Publication |
|---|---|---|---|
| **1** | 3-3-1 | 1242 | Zacharaki et al., 2022 |
| **2** | 140-2-1 | 1179 | Zacharaki et al., 2022 |
| **3** | 1-3-2 | 1008 | Zacharaki et al., 2022 |
| **4** | 57-1-2 | 992 | Zacharaki et al., 2022 |
| **5** | 228-1-2 | 946 | Zacharaki et al., 2022 |
| **6** | 160-1 | 907 | Zacharaki et al., 2022 |
| **7** | 75-2-1 | 904 | Zacharaki et al., 2022 |
| **8** | 175-2-1 | 880 | Zacharaki et al., 2022 |
| **9** | 233-14-1 | 880 | Zacharaki et al., 2022 |
| **10** | 243-5-1 | 863 | Zacharaki et al., 2022 |
| **11** | 92-2-1 | 855 | Zacharaki et al., 2022 |
| **12** | 130-1-1 | 854 | Zacharaki et al., 2022 |
| **13** | 8-1-1 | 853 | Zacharaki et al., 2022 |
| **14** | 57-2-1 | 845 | Zacharaki et al., 2022 |
| **15** | 79-5-2 | 842 | Zacharaki et al., 2022 |
| **16** | 183-1-1 | 819 | Zacharaki et al., 2022 |
| **17** | 255-5-1 | 816 | Zacharaki et al., 2022 |
| **18** | 199-1-1 | 809 | Zacharaki et al., 2022 |
| **19** | 72-1-2 | 790 | Zacharaki et al., 2022 |
| **20** | 79-4-1 | 785 | Zacharaki et al., 2022 |
| **21** | 144-1-1 | 781 | Zacharaki et al., 2022 |
| **22** | 225-5-1 | 777 | Zacharaki et al., 2022 |
| **23** | 144-2-1 | 774 | Zacharaki et al., 2022 |
| **24** | 125-6-1 | 773 | Zacharaki et al., 2022 |
| **25** | 58-6-2 | 757 | Zacharaki et al., 2022 |
| **26** | 158-5-2 | 743 | Zacharaki et al., 2022 |
| **27** | 103-5-1 | 714 | Zacharaki et al., 2022 |
| **28** | 32-7-1 | 710 | Zacharaki et al., 2022 |
| **29** | 48-2-2 | 710 | Zacharaki et al., 2022 |
| **30** | 92-3-2 | 694 | Zacharaki et al., 2022 |
| **31** | 54-4-1 | 693 | Zacharaki et al., 2022 |
| **32** | 48-1-2 | 683 | Zacharaki et al., 2022 |
| **33** | 170-1-1 | 678 | Zacharaki et al., 2022 |
| **34** | 236-10-1 | 659 | Zacharaki et al., 2022 |
| **35** | 91-1-21-2 | 653 | Zacharaki et al., 2022 |
| **36** | 131-13-1 | 649 | Zacharaki et al., 2022 |
| **37** | 101-4-2 | 645 | Zacharaki et al., 2022 |
| **38** | 171-2-1 | 645 | Zacharaki et al., 2022 |
| **39** | 171-1-1 | 637 | Zacharaki et al., 2022 |
| **40** | 199-6-2 | 634 | Zacharaki et al., 2022 |
| **41** | 184-1-2 | 620 | Zacharaki et al., 2022 |
| **42** | 244-2-2 | 617 | Zacharaki et al., 2022 |
| **43** | 155-3-1 | 613 | Zacharaki et al., 2022 |
| **44** | 154-1-1 | 597 | Zacharaki et al., 2022 |
| **45** | 103-2-2 | 595 | Zacharaki et al., 2022 |
| **46** | 101-3-1 | 594 | Zacharaki et al., 2022 |
| **47** | 155-2-1 | 542 | Zacharaki et al., 2022 |
| **48** | 131-29-2 | 526 | Zacharaki et al., 2022 |
| **49** | 225-1-1 | 519 | Zacharaki et al., 2022 |
| **50** | 105-1-2 | 489 | Zacharaki et al., 2022 |
| **51** | 163-5-2 | 481 | Zacharaki et al., 2022 |
| **52** | 232-2-1 | 474 | Zacharaki et al., 2022 |
| **53** | 196-1-2 | 453 | Zacharaki et al., 2022 |
| **54** | 212-2-2 | 383 | Zacharaki et al., 2022 |
| **55** | 180-3-1 | 343 | Zacharaki et al., 2022 |
| **56** | 164-9-1 | 328 | Zacharaki et al., 2022 |
| **57** | 180-1-1 | 282 | Zacharaki et al., 2022 |
| **58** | 50-1-2 | 33 | Zacharaki et al., 2022 |
| **59** | 292-2-1 | 31 | Zacharaki et al., 2022 |
| **60** | 75-1-1 | 25 | Zacharaki et al., 2022 |
| **61** | 233-13-1 | 24 | Zacharaki et al., 2022 |
| **62** | 192-1-2 | 22 | Zacharaki et al., 2022 |
| **63** | 292-1-1 | 21 | Zacharaki et al., 2022 |
| **64** | 192-2-2 | 20 | Zacharaki et al., 2022 |
| **65** | 230-2-2 | 17 | Zacharaki et al., 2022 |
| **66** | 49-5-3 | 1523 | this publication |
| **67** | 132-1-2 | 1372 | this publication |
| **68** | 219-1-2 | 1100 | this publication |
| **69** | 91-1-21 | 980 | this publication |
| **70** | 106-2-1 | 731 | this publication |
| **71** | 232-2-2 | 716 | this publication |
| **72** | 292-1-2 | 74 | this publication |
| **73** | 34-8-1 | 74 | this publication |
| **74** | 192-1-1 | 64 | this publication |
| **75** | 50-1-1 | 82 | this publication |
| **76** | 230-2 | 63 | this publication |
| **77** | 278-1-1 | 62 | this publication |
| **78** | 30-34 | 941 | this publication |
| **79** | 107-2 | 1032 | this publication |
| **80** | 42-9 | 418 | this publication |
| **81** | 128-1 | 104 | this publication |
| **82** | 30-23 | 220 | this publication |
| **83** | 55-21 | 1035 | this publication |
| **84** | 55-6 | 905 | this publication |
| **85** | 55-15 | 371 | this publication |
| **86** | 11-7 | 833 | this publication |
| **87** | 77-3 | 1246 | this publication |
| **88** | 2-1 | 1240 | this publication |
| **89** | 160-1-b | 916 | this publication |
| **90** | 161-1 | 685 | this publication |
| **91** | 271-1 | 1384 | this publication |
| **92** | 41-18 | 655 | this publication |

**Table S2. Number of SNPs identified in EMS suppressor lines carrying mutations in *HUA2*.**

| Line ID | DNA source | Number SNPs |
|---|---|---|
| 8-1-1 | Individual suppressor mutant | 853 |
| 30-34 | Individual suppressor mutant | 941 |
| 57-2-1 | Individual suppressor mutant | 845 |
| 233-14-1 | Individual suppressor mutant | 880 |
| 164-9-1 | Individual suppressor mutant | 328 |

**Table S3. EMS suppressor lines bearing non-synonymous mutations in *HUA2*.**

| Gene isoform | Number of line(s) SNP is present | Line(s) ID | Chromosomal SNP position | Reference base | Alternative base | Feature | Codon position in gene | Length of CDS | Position of SNP in the CDS | Position of SNP in codon | Non-synonymous/ Synonymous | Reference amino acid | Alternative amino acid | Degeneracy |
|---|---|---|---|---|---|---|---|---|---|---|---|---|---|---|
| AT5G23150.1 | 3 | 8-1-1<br>30-34<br>57-2-1 | 7790473 | G | A | CDS | 4639 | 4179 | 2947 | 1 | Nonsyn | A | T | Gcg |
| AT5G23150.1 | 1 | 233-14-1 | 7788166 | C | T | CDS | 2332 | 4179 | 1363 | 1 | Nonsyn | P | S | Cca |
| AT5G23150.1 | 1 | 164-9-1 | 7790099 | C | T | CDS | 4265 | 4179 | 2704 | 1 | Nonsyn | R | C | Cgt |

**Table S6. Expression of flowering time genes in *hua2-4 tps1-2 GVG::TPS1* and *tps1-2 GVG::TPS1*.**

| Gene_ID | start | end | strand | *tps1-2 GVG::TPS1 hua2-4* | | | *tps1-2 GVG::TPS1* | | |
|---|---|---|---|---|---|---|---|---|---|
| Replicate | | | | 1 | 2 | 3 | 1 | 2 | 3 |
| AT4G24540(*AGL24*) | 12670965 | 12674072 | - | 23,124781 | 25,085474 | 23,208811 | 11,087811 | 13,027147 | 10,491028 |
| AT2G45660(*SOC1*) | 18807538 | 18811047 | - | 8,265232 | 10,200581 | 12,231363 | 4,341604 | 4,784088 | 6,764344 |
| AT5G61850(*LFY*) | 24844295 | 24846933 | + | 0 | 0 | 0 | 0 | 0 | 0 |
| AT5G10140(*FLC*) | 3173497 | 3179448 | - | 0 | 0 | 0 | 0 | 0 | 0 |
| AT5G15840(*CO*) | 5171182 | 5172758 | - | 0,013072 | 0,140295 | 0,169611 | 0,194254 | 0 | 0,017146 |
| AT1G65480(*FT*) | 24331428 | 24333934 | + | 0,419172 | 0,094804 | 0,110622 | 0,11757 | 0,109185 | 0 |
| AT4G20370(*TSF*) | 11000771 | 11002996 | - | 0 | 0,059687 | 0,593648 | 0,125811 | 0 | 0 |
| AT3G54990(*SMZ*) | 20373718 | 20376522 | - | 0 | 0,002427 | 0,005711 | 0,00271 | 0,00165 | 0,009862 |
| AT2G33810(*SPL3*) | 14305001 | 14306072 | + | 4,384162 | 4,665609 | 7,363255 | 5,170671 | 4,712061 | 3,068198 |
| AT1G53160(*SPL4*) | 19806419 | 19807608 | + | 3,940252 | 5,441206 | 6,007432 | 5,306567 | 5,117388 | 3,515887 |
| AT3G15270(*SPL5*) | 5140365 | 5141348 | - | 0,289313 | 0,464 | 0,67997 | 0,533319 | 0,719163 | 0,33364 |
| AT2G28550(*TOE1*) | 12225842 | 12228543 | - | 4,908707 | 4,502523 | 6,183015 | 7,100536 | 7,014573 | 7,059748 |
| AT5G60120(*TOE2*) | 24207786 | 24211724 | + | 0,373802 | 0,578502 | 0,191514 | 0,318693 | 0,131952 | 0,211047 |
| AT5G67180(*TOE3*) | 26801949 | 26804249 | - | 1,491339 | 0,927218 | 1,287269 | 0,731689 | 1,146463 | 1,783746 |
| AT2G39250(*SNZ*) | 16388886 | 16391073 | - | 0,723291 | 0,935247 | 0,324612 | 0,66105 | 0,179 | 0,620404 |

**Table S7. List of oligonucleotides used in this study.**

| **Purpose** | ***Target*** | **Sequence (5' => 3')** |
|---|---|---|
| Genotyping *ft-10* | *ft-10* | AGGGTTGCTAGGACTTGGAACA |
| Genotyping *ft-10* | *ft-10* | CCCATTTGGACGTGAATGTAGACAC |
| Genotyping *ft-10* | *T-DNA* border | GGTGGAGAAGACCTCAGGAAC |
| Genotyping *hua2-4* | *hua2-4* | TCTATCAGAGCCACCTGCTTC |
| Genotyping *hua2-4* | *hua2-4* | TTACTCGGTCAGATTCCATGG |
| Genotyping *hua2-4* | *T-DNA* border | ATTTTGCCGATTTCGGAAC |
| Genotyping *soc1* | *soc1* | TGTGTGCAAGGGAAATTAACTAAAGAAGAAGAT |
| Genotyping *soc1* | *soc1* | TTAGTATGCCTCAGATAACGATCTATGGTAT |
| Genotyping *soc1* | *T-DNA* border | ATTTTGCCGATTTCGGAAC |
| Genotyping *flc* | *flc* | AGCCAAGAAGACCGAACTCA |
| Genotyping *flc* | *flc* | TTTGTCCAGCAGGTGACATC |
| Genotyping and sequencing EMS lines (233-14-1) | *hua2* | TGGAGCATGCTACCTCTCCT |
| Genotyping and sequencing EMS lines (233-14-1) | *hua2* | TCCTGCAACGTGCTTGTTAG |
| Genotyping and sequencing EMS lines (164-9-1) | *hua2* | TCCTGAAGTTGTGGCTTGAA |
| Genotyping and sequencing EMS lines (164-9-1) | *hua2* | TTCATGATCGACATCCTCCA |
| Genotyping and sequencing EMS lines (8-1-1, 30-34, 57-2-1) | *hua2* | TTTTTCTGCAGCAATGCAAC |
| Genotyping and sequencing EMS lines (8-1-1, 30-34, 57-2-1) | *hua2* | AGGTGGAGGTGAAAGTGGTG |
| Genotyping *tps1-2* | *tps1-2* | GACACTTGGTTTCTTGATATGTCCTG |
| Genotyping *tps1-2* | *tps1-2* | GCTGTCTTGGATACTGAACCAGT |
| Genotyping *tps1-2* | *tps1-2* transposon | GAGCGTCGGTCCCCACACTTCTATAC |
| RT-qPCR | *TUB2* | GAGCCTTACAACGCTACTCTGTCTGTC |
| RT-qPCR | *TUB2* | ACACCAGACATAGTAGCAGAAATCAAG |
| RT-qPCR | *UBI* | CACACTCCACTTGGTCTTGCGT |
| RT-qPCR | *UBI* | TGGTCTTTCCGGTGAGAGTCTTCA |
| RT-qPCR | *TPS1* | GAAACTCAAGACGTCCTTCACCAG |
| RT-qPCR | *TPS1* | TCTAGCATTGGTGCGAGTACGAC |
| RT-qPCR | *SOC1* | TTGAGCAGCTCAAGCAAAAGGA |
| RT-qPCR | *SOC1* | TCCCCACTTTTCAGAGAGCTTCTC |
| RT-qPCR | *FT* | CCCTGCTACAACTGGAACAAC |
| RT-qPCR | *FT* | CACCCTGGTGCATACACTG |
| RT-qPCR | *FLC* | AGCCAAGAAGACCGAACTCA |
| RT-qPCR | *FLC* | TTTGTCCAGCAGGTGACATC |
